## Supplemental File 1-5 for "Modular DNA Barcoding of Nanobodies Enables Multiplexed *in situ* Protein Imaging and High-throughput Biomolecule Detection"

### Supplementary File 1. Detailed compound information in HTS-based BLISA.

| drug# | Name | CAS | MolWt | Vector | Cat.No. |
|---|---|---|---|---|---|
| **D1** | Flubendazole | 31430-15-6 | 313.29 | TopScience | TS0009 |
| **D2** | Disulfiram | 97-77-8 | 296.54 | TopScience | TS0050 |
| **D3** | RISPERIDONE | 106266-06-2 | 410.48 | TopScience | TS0052 |
| **D4** | Chlorpromazine hydrochloride | 69-09-0 | 355.32 | TopScience | TS0067 |
| **D5** | PHENFORMIN HYDROCHLORIDE | 834-28-6 | 241.72 | TopScience | TS0099 |
| **D6** | Paroxetine HCl | 78246-49-8 | 365.83 | TopScience | TS0131 |
| **D7** | CLOMIPRAMINE HYDROCHLORIDE | 17321-77-6 | 351.31 | TopScience | TS0183 |
| **D8** | HALOPERIDOL | 52-86-8 | 375.8744 | TopScience | TS0395 |
| **D9** | FLUPHENAZINE HYDROCHLORIDE | 146-56-5 | 510.44 | TopScience | TS0401 |
| **D10** | CARBAMAZEPINE | 298-46-4 | 236.27 | TopScience | TS0450 |
| **D11** | CLOZAPINE | 5786-21-0 | 326.82 | TopScience | TS0471 |
| **D12** | Levetiracetam | 102767-28-2 | 170.21 | TopScience | TS0506 |
| **D13** | Citalopram HBr | 59729-32-7 | 405.31 | TopScience | TS0567 |
| **D14** | DOXEPIN HYDROCHLORIDE | 1229-29-4 | 315.84 | TopScience | TS0619 |
| **D15** | TOPOTECAN HYDROCHLORIDE | 119413-54-6 | 457.9178 | TopScience | TS0694 |
| **D16** | NISOLDIPINE | 63675-72-9 | 388.41 | TopScience | TS0719 |
| **D17** | QUETIAPINE | 111974-69-7 | 383.51 | TopScience | TS0734 |
| **D18** | Epirubicin hydrochloride | 56390-09-1 | 579.9802 | TopScience | TS0759 |
| **D19** | THIORIDAZINE HYDROCHLORIDE | 130-61-0 | 407.0437 | TopScience | TS0777 |
| **D20** | VINCRISTINE SULFATE | 2068-78-2 | 923.04 | TopScience | TS0862 |
| **D21** | CLOFAZIMINE | 2030-63-9 | 473.4 | TopScience | TS0889 |
| **D22** | ARIPIPRAZOLE | 129722-12-9 | 448.39 | TopScience | TS0896 |
| **D23** | ITRACONAZOLE | 84625-61-6 | 705.63 | TopScience | TS0898 |
| **D24** | Sodium 2-propylpentanoate | 1069-66-5 | 166.2 | TopScience | TS0907 |
| **D25** | AMOXAPINE | 14028-44-5 | 313.7896 | TopScience | TS0999 |
| **D26** | OLANZAPINE | 132539-06-1 | 312.4398 | TopScience | TS1002 |
| **D27** | SERTRALINE HYDROCHLORIDE | 79559-97-0 | 342.69 | TopScience | TS1008 |
| **D28** | TRIFLUOPERAZINE HYDROCHLORIDE | 440-17-5 | 480.4267 | TopScience | TS1038 |
| **D29** | PERPHENAZINE | 58-39-9 | 403.97 | TopScience | TS1113 |
| **D30** | FLUVOXAMINE MALEATE | 61718-82-9 | 434.41 | TopScience | TS1130 |
| **D31** | AMITRIPTYLINE HYDROCHLORIDE | 549-18-8 | 313.86 | TopScience | TS1250 |
| **D32** | Docetaxel | 114977-28-5 | 807.88 | TopScience | T1034 |
| **D33** | Vandetanib | 443913-73-3 | 475.31 | TopScience | T1656 |
| **D34** | Pitavastatin (Calcium) | 147526-32-7 | 420.45 | MCE | HY-B0144 |
| **D35** | Asenapine (maleate) | 85650-56-2 | 401.84 | MCE | HY-11100 |
| **D36** | Vinorelbine (ditartrate) | 125317-39-7 | 1079.11 | MCE | HY-12053A |
| **D37** | Doxorubicin (hydrochloride) | 25316-40-9 | 579.98 | MCE | HY-15142 |
| **D38** | OSI-420 | 183320-51-6 | 415.87 | MCE | HY-13256 |

| | | | | | |
|---|---|---|---|---|---|
| **D39** | Lurasidone (Hydrochloride) | 367514-88-3 | 529.14 | MCE | HY-B0032 |
| **D40** | Irinotecan (hydrochloride) | 100286-90-6 | 623.14 | MCE | HY-16562A |
| **D41** | Escitalopram (oxalate) | 219861-08-2 | 414.43 | MCE | HY-14258A |
| **D42** | Homoharringtonine | 26833-87-4 | 545.62 | MCE | HY-14944 |
| **D43** | Fluoxetine (hydrochloride) | 56296-78-7 | 345.79 | MCE | HY-B0102A |
| **D44** | PD184352 (CI-1040) | 212631-79-3 | 478.67 | Selleck | S1020 |
| **D45** | Ridaforolimus (Deforolimus, MK-8669) | 572924-54-0 | 990.21 | Selleck | S1022 |
| **D46** | Nilotinib (AMN-107) | 641571-10-0 | 529.52 | Selleck | S1033 |
| **D47** | Sunitinib Malate | 341031-54-7 | 532.56 | Selleck | S1042 |
| **D48** | Masitinib (AB1010) | 790299-79-5 | 498.64 | Selleck | S1064 |
| **D49** | ZSTK474 | 475110-96-4 | 417.41 | Selleck | S1072 |
| **D50** | MLN8054 | 869363-13-3 | 476.86 | Selleck | S1100 |
| **D51** | LY294002 | 154447-36-6 | 307.34 | Selleck | S1105 |
| **D52** | OSU-03012 (AR-12) | 742112-33-0 | 460.45 | Selleck | S1106 |
| **D53** | BX-912 | 702674-56-4 | 471.35 | Selleck | S1275 |
| **D54** | Pelitinib (EKB-569) | 257933-82-7 | 467.92 | Selleck | S1392 |
| **D55** | Aurora A Inhibitor I | 1158838-45-9 | 588.07 | Selleck | S1451 |
| **D56** | HMN-214 | 173529-46-9 | 424.47 | Selleck | S1485 |
| **D57** | CP-673451 | 343787-29-1 | 417.5 | Selleck | S1536 |
| **D58** | BS-181 HCl | 1397219-81-6 | 416.99 | Selleck | S1572 |
| **D59** | Tie2 kinase inhibitor | 948557-43-5 | 439.53 | Selleck | S1577 |
| **D60** | BMS-265246 | 582315-72-8 | 345.34 | Selleck | S2014 |
| **D61** | GSK461364 | 929095-18-1 | 543.6 | Selleck | S2193 |
| **D62** | Chrysophanic Acid | 481-74-3 | 254.24 | Selleck | S2406 |
| **D63** | Phenformin HCl | 834-28-6 | 241.72 | Selleck | S2542 |
| **D64** | Trametinib (GSK1120212) | 871700-17-3 | 615.39 | Selleck | S2673 |
| **D65** | CHIR-124 | 405168-58-3 | 419.91 | Selleck | S2683 |
| **D66** | KX2-391 | 897016-82-9 | 431.53 | Selleck | S2700 |
| **D67** | ZM 336372 | 208260-29-1 | 389.45 | Selleck | S2720 |
| **D68** | BGT226 (NVP-BGT226) | 1245537-68-1 | 650.6 | Selleck | S2749 |
| **D69** | CEP-33779 | 1257704-57-6 | 462.57 | Selleck | S2806 |
| **D70** | GDC-0068 | 1001264-89-6 | 458 | Selleck | S2808 |
| **D71** | TAE226 (NVP-TAE226) | 761437-28-9 | 468.94 | Selleck | S2820 |
| **D72** | Semaxanib (SU5416) | 194413-58-6 | 238.28 | Selleck | S2845 |
| **D73** | JNK-IN-8 | 1410880-22-6 | 507.59 | Selleck | S4901 |
| **D74** | 10058-F4 | 403811-55-2 | 249.35 | Selleck | S7153 |
| **D75** | LY2835219 | 1231930-82-7 | 602.7 | Selleck | S7158 |
| **D76** | SSR128129E | 848318-25-2 | 346.31 | Selleck | S7167 |
| **D77** | CNX-774 | 1202759-32-7 | 499.5 | Selleck | S7257 |
| **D78** | PFK15 | 4382-63-2 | 260.29 | Selleck | S7289 |
| **D79** | TAK-632 | 1228591-30-7 | 554.52 | Selleck | S7291 |
| **D80** | AZD9291 | 1421373-65-0 | 499.61 | Selleck | S7297 |
| **D81** | KN-62 | 127191-97-3 | 721.84 | Selleck | S7422 |
| **D82** | GNE-7915 | 1351761-44-8 | 443.4 | Selleck | S7528 |

| | | | | | |
|---|---|---|---|---|---|
| **D83** | IM-12 | 1129669-05-1 | 377.41 | Selleck | S7566 |
| **D84** | SMI-4a | 438190-29-5 | 273.23 | Selleck | S8005 |
| **D85** | NSC 23766 | 1177865-17-6 | 530.96 | Selleck | S8031 |
| **D86** | Pacritinib (SB1518) | 937272-79-2 | 472.58 | Selleck | S8057 |
| **D87** | LY2228820 | 862507-23-1 | 612.74 | Selleck | S1494 |
| **D88** | Idarubicin hydrochloride | 57852-57-0 | 533.954 | TopScience | T6010 |
| **D89** | Imipramine (hydrochloride) | 113-52-0 | 316.8682 | MCE | HY-B1490 |
| **D90** | NU7441 | 503468-95-9 | 413.48826 | Selleck | S2638 |

**Supplementary File 2. Antibodies and Nb-DNA oligos (or Nb-SS-DNA oligos).**

| **Epitope** | **Host** | **Nb-DNA oligos (or Nb-SS-DNA oligos)** |
|---|---|---|
| PDGFRα | Rabbit | TP897-B1 I1 |
| KRT14 | Rabbit | TP897-B3 I1 |
| DCT | Rabbit | TP897-B4 I1 |
| CD31 | Mouse (IgG1) | TP1107-B9 I1 |
| αSMA | Rabbit | TP897-B10 I1 |
| CD45 | Mouse (IgG1) | TP1107-B13 I1 |
| MAP2 | Rabbit | TP897-B1 I1 |
| NeuN | Rabbit | TP897-B1 I1, TP897-SS-B15 I1 |
| TPH2 | Rabbit | TP897-B5 I1, TP897-SS-B5 I1 |
| TH | Rabbit | TP897-B1 I1, TP897-SS-B14 I1 |
| nNOS | Rabbit | TP897-B5 I1, TP897-SS-B10 I1 |
| GFAP | Rabbit | TP897-B5 I1, TP897-SS-B13 I1 |
| Iba1 | Rabbit | TP897-B4 I1, TP897-SS-B13 I1 |
| DDC | Rabbit | TP897-B4 I1, TP897-SS-B14 I1 |
| NF-H | Rabbit | TP897-B4 I1, TP897-SS-B17 I1 |
| TMEM119 | Rabbit | TP897-B1 I1, TP897-SS-B15 I1 |
| GABA | Rabbit | TP897-B1 I1, TP897-SS-B2 I1 |
| Orexin A | Rabbit | TP897-B1 I1, TP897-SS-B9 I1 |
| 5-HT | Rabbit | TP897-B1 I1, TP897-SS-B4 I1 |
| α-tubulin | Mouse (IgG1) | TP1107-SS-qbc. 1, TP1107-SS-sbc. 1 |
| GFP | Rabbit | TP897-SS-qbc. 2, TP897-SS -sbc. 2 |
| mCherry | Rabbit | TP897-SS-qbc. 3, TP897-SS-sbc. 3 |
| Human IgG | Rabbit | TP897-SS-qbc. 2, TP897-SS-sbc. 4 |
| HBsAg (capture) | Goat | |
| HBsAg (detection) | Mouse (IgG1) | TP1107-SS-qbc. 3, TP1107-SS-sbc. 3 |
| HBeAg (capture) | Mouse (IgG2a) | |
| HBeAg (detection) | Mouse (IgG1) | TP1107-SS-qbc. 2, TP1107-SS-sbc. 2 |
| phospho-p38α (T180/Y182) (capture) | Mouse | |
| phospho-p38α (T180/Y182) (detection) | Rabbit | TP897-SS-qbc. 2, TP897-SS-sbc. 2 |
| phospho-ERK1 (T202/Y204)/ERK2 (T185/Y187) (capture) | Mouse | |
| phospho-ERK1 (T202/Y204)/ERK2 (T185/Y187) (detection) | Rabbit | TP897-SS-qbc. 3, TP897-SS-sbc. 3 |
| phospho-JNK Pan Specific (capture) | Mouse | |
| phospho-JNK Pan Specific (detection) | Rabbit | TP897-SS-qbc. 4, TP897-SS-sbc. 4 |
| phospho-AMPKα1 (T183) (capture) | Goat | |
| phospho-AMPKα1 (T183) (detection) | Rabbit | TP897-SS-qbc. 5, TP897-SS-sbc. 5 |
| phospho-CREB (S133) (capture) | Goat | |
| phospho-CREB (S133) (detection) | Rabbit | TP897-SS-qbc. 6, TP897-SS-sbc. 6 |
| phospho-Src (Y419) (capture) | Goat | |
| phospho-Src (Y419) (detection) | Rabbit | TP897-SS-qbc. 7, TP897-SS-sbc. 7 |
| phospho-Akt (S473) (capture) | Rabbit | |

| | | |
|---|---|---|
| phospho-Akt (S473) (detection) | Rabbit | TP1107-SS-qbc. 1, TP1107-SS-sbc. 1 |

**Supplementary File 3. Detailed sequences and modifications of DNA oligos.**

| Name | Sequence (5' to 3') | Modifications |
|---|---|---|
| **B1 I1** | ATATAGCATTCTTTCTTGAGGAGGGCAGCAAACGGGAAGAG | 5' DBCO or 5' Amino C6 |
| **B2 I1** | ATATAAGCTCAGTCCATCCTCGTAAATCCTCATCAATCATC | 5' DBCO or 5' Amino C6 |
| **B3 I1** | ATATAAAAGTCTAATCCGTCCCTGCCTCTATATCTCCACTC | 5' DBCO or 5' Amino C6 |
| **B4 I1** | ATATACACATTTACAGACCTCAACCTACCTCCAACTCTCAC | 5' DBCO or 5' Amino C6 |
| **B5 I1** | ATATACACTTCATATCACTCACTCCCAATCTCTATCTACCC | 5' DBCO or 5' Amino C6 |
| **B9 I1** | ATATACACGTATCTACTCCACTCTCAGCACACTCCCAACCC | 5' DBCO or 5' Amino C6 |
| **B10 I1** | ATATACCTCAAGATACTCCTCTACCTACTCGACTACCCTAG | 5' DBCO or 5' Amino C6 |
| **B13 I1** | ATATAAGGTAACGCCTTCCTGCTTTATGCTCAACATACAAC | 5' DBCO or 5' Amino C6 |
| **B14 I1** | ATATAAATGTCAATAGCGAGCGACCCTATATTCTGCACAG | 5' DBCO or 5' Amino C6 |
| **B15 I1** | ATATACAGATTAACACACCACAAGGTATCTCGAACACTCTC | 5' DBCO or 5' Amino C6 |
| **B17 I1** | ATATACGATTGTTTGTTGTGGACGCATGCTAATCGGATGAG | 5' DBCO or 5' Amino C6 |
| **B1 Amplifier H1** | CGTAAAGGAAGACTCTTCCCGTTTGCTGCCCTCCTCGCATTCTTTCTT GAGGAGGGCAGCAAACGGGAAGAG | 5' Alexa Fluor 546 or 5' Cy3 |
| **B1 Amplifier H2** | GAGGAGGGCAGCAAACGGGAAGAGTCTTCCTTTACGCTCTTCCCGTT TGCTGCCCTCCTCAAGAAAGAATGC | 3' Alexa Fluor 546 or 3' Cy3 |
| **B2 Amplifier H1** | GGCGGTTTACTGGATGATTGATGAGGATTTACGAGGAGCTCAGTCCA TCCTCGTAAATCCTCATCAATCATC | 5' Alexa Fluor 594 |
| **B2 Amplifier H2** | CCTCGTAAATCCTCATCAATCATCCAGTAAACCGCCGATGATTGATG AGGATTTACGAGGATGGACTGAGCT | 3' Alexa Fluor 594 |
| **B3 Amplifier H1** | CGGGTTAAAGTTGAGTGGAGATATAGAGGCAGGGACAAAGTCTAAT CCGTCCCTGCCTCTATATCTCCACTC | 5' Alexa Fluor 488 |
| **B3 Amplifier H2** | GTCCCTGCCTCTATATCTCCACTCAACTTTAACCCGGAGTGGAGATAT AGAGGCAGGGACGGATTAGACTTT | 3' Alexa Fluor 488 |
| **B4 Amplifier H1** | GAAGCGAATATGGTGAGAGTTGGAGGTAGGTTGAGGCACATTTACA GACCTCAACCTACCTCCAACTCTCAC | 5' Alexa Fluor 647 or 5' Cy5 |
| **B4 Amplifier H2** | CCTCAACCTACCTCCAACTCTCACCATATTCGCTTCGTGAGAGTTGGA GGTAGGTTGAGGTCTGTAAATGTG | 3' Alexa Fluor 647 or 3' Cy5 |
| **B5 Amplifier H1** | ATTGGATTTGTAGGGTAGATAGAGATTGGGAGTGAGCACTTCATATC ACTCACTCCCAATCTCTATCTACCC | 5' Alexa Fluor 488 |
| **B5 Amplifier H2** | CTCACTCCCAATCTCTATCTACCCTACAAATCCAATGGGTAGATAGA GATTGGGAGTGAGTGATATGAAGTG | 3' Alexa Fluor 488 |
| **B9 Amplifier H1** | CCACTCTCAGCACACTCCCAACCCTACTACAAGCTCGGGTTGGGAGT GTGCTGAGAGTGGAGTAGATACGTG | 5' Alexa Fluor 647 |
| **B9 Amplifier H2** | GAGCTTGTAGTAGGGTTGGGAGTGTGCTGAGAGTGGCACGTATCTAC TCCACTCTCAGCACACTCCCAACCC | 3' Alexa Fluor 647 |

| | | |
|---|---|---|
| **B10 Amplifier H1** | CCTCTACCTACTCGACTACCCTAGCCGTAACTTCACCTAGGGTAGTCGAGTAGGTAGAGGAGTATCTTGAGG | 5' Alexa Fluor 488 |
| **B10 Amplifier H2** | GTGAAGTTACGGCTAGGGTAGTCGAGTAGGTAGAGGCCTCAAGATACTCCTCTACCTACTCGACTACCCTAG | 3' Alexa Fluor 488 |
| **B13 Amplifier H1** | CCTGCTTTATGCTCAACATACAACCAGAAATGCGGCGTTGTATGTTGAGCATAAAGCAGGAAGGCGTTACCT | 5' Alexa Fluor 647 |
| **B13 Amplifier H2** | GCCGCATTTCTGGTTGTATGTTGAGCATAAAGCAGGAGGTAACGCCTTCCTGCTTTATGCTCAACATACAAC | 3' Alexa Fluor 647 |
| **B14 Amplifier H1** | GAGCGACCCTATATTTCTGCACAGAAGTTATACCGGCTGTGCAGAAATATAGGGTCGCTCGCTATTGACATT | 5' Alexa Fluor 488 |
| **B14 Amplifier H2** | CCGGTATAACTTCTGTGCAGAAATATAGGGTCGCTCAATGTCAATAGCGAGCGACCCTATATTTCTGCACAG | 3' Alexa Fluor 488 |
| **B15 Amplifier H1** | CCACAAGGTATCTCGAACACTCTCCAAATTGGCTACGAGAGTGTTCGAGATACCTTGTGGTGTGTTAATCTG | 5' Alexa Fluor 546 |
| **B15 Amplifier H2** | GTAGCCAATTTGGAGAGTGTTCGAGATACCTTGTGGCAGATTAACACACCACAAGGTATCTCGAACACTCTC | 3' Alexa Fluor 546 |
| **B17 Amplifier H1** | GTGGACACCTGCTAATCGGATGAGTGTTCGTTATCGCTCATCCGATTAGCAGGTGTCCACAACAAACAATCG | 5' Alexa Fluor 647 |
| **B17 Amplifier H2** | CGATAACGAACACTCATCCGATTAGCAGGTGTCCACCGATTGTTTGTTGTGGACACCTGCTAATCGGATGAG | 3' Alexa Fluor 647 |
| **qbc. 1 (for qPCR)** | TCTTGTGGAAAGGACGAAACACGTGATNNNNNNNNNNNNNNNNGTCTGGAGCATGCGCTTTAG | 5' Amino C6 |
| **qbc. 2 (for qPCR)** | TACACGACGCTCTTCCGATCTCGTGATNNNNNNNNNNNNNNNNTTGAAAAAGTGGCACCGAGT | 5' Amino C6 |
| **qbc. 3 (for qPCR)** | ACACGTCTGAACTCCAGTCACCGTGATNNNNNNNNNNNNNNNNCGTATGCCGTCTTCTGCTTG | 5' Amino C6 |
| **qbc. 4 (for qPCR)** | GACAGTTCGAGTTTGAAGCGCGTGATNNNNNNNNNNNNNNNNCTAGACGTGGGAGTGCATACT | 5' Amino C6 |
| **qbc. 5 (for qPCR)** | GAAAGATCTGGCTGCCATGCCGTGATNNNNNNNNNNNNNNNNTCGCAAACCTGGTTGGAATCA | 5' Amino C6 |
| **qbc. 6 (for qPCR)** | AGATGACGTCGATTGTTGGTCGTGATNNNNNNNNNNNNNNNNCATGGAGGTTGTGTCACCGTA | 5' Amino C6 |
| **qbc. 7 (for qPCR)** | TCAGGTGCATAGGAGTCAGCCGTGATNNNNNNNNNNNNNNNNATGCTGTCAGTTCATGGCTCC | 5' Amino C6 |
| **sbc. 1 (for sequencing)** | TCTTGTGGAAAGGACGAAACA**CGTGAT**NNNNNNNNNNNNNNNNGTCTGGAGCATGCGCTTTAG | 5' Amino C6 |
| **sbc. 2 (for sequencing)** | TCTTGTGGAAAGGACGAAACA**ATCACG**NNNNNNNNNNNNNNNNGTCTGGAGCATGCGCTTTAG | 5' Amino C6 |
| **sbc. 3 (for sequencing)** | TCTTGTGGAAAGGACGAAACA**CGATGT**NNNNNNNNNNNNNNNNGTCTGGAGCATGCGCTTTAG | 5' Amino C6 |
| **sbc. 4 (for sequencing)** | TCTTGTGGAAAGGACGAAACA**TTAGGC**NNNNNNNNNNNNNNNNGTCTGGAGCATGCGCTTTAG | 5' Amino C6 |
| **sbc. 5 (for sequencing)** | TCTTGTGGAAAGGACGAAACA**TGACCA**NNNNNNNNNNNNNNNNGTCTGGAGCATGCGCTTTAG | 5' Amino C6 |
| **sbc. 6 (for sequencing)** | TCTTGTGGAAAGGACGAAACA**ACAGTG**NNNNNNNNNNNNNNNNGTCTGGAGCATGCGCTTTAG | 5' Amino C6 |
| **sbc. 7 (for sequencing)** | TCTTGTGGAAAGGACGAAACA**GCCAAT**NNNNNNNNNNNNNNNNGTCTGGAGCATGCGCTTTAG | 5' Amino C6 |

| | |
|---|---|
| **Norm DNA (for sequencing)** | TCTTGTGGAAAGGACGAAACA**GCCAAT**NNNNNNNNNNNNNNNNGTCTGGAGCATGCGCTTTAG |

The sequences of Ab bc/norm bc were underlined.

**Supplementary File 4. Amino acid sequences of the constructs used in this study.**

pET28a-His$_6$-Ubiquitin-OaAEP1 (C247A):

MGSSHHHHHHSSGENLYFQGRPMQIFVKTLTGKTITLEVEPSDTIENVKAKIQDKEGIPPDQQRLIFAGKQLEDGRTLS
DYNIQKESTLHLVLRLRGGARDGDYLHLPSEVSRFFRPQETNDDHGEDSVGTRWAVLIAGSKGYANYRHQAGVCHA
YQILKRGGLKDENIVVFMYDDIAYNESNPRPGVIINSPHGSDVYAGVPKDYTGEEVNAKNFLAAILGNKSAITGGSG
KVVDSGPNDHIFIYYTDHGAAGVIGMPSKPYLYADELNDALKKKHASGTYKSLVFYLEACESGSMFEGILPEDLNIY
ALTSTNTTESSWAYYCPAQENPPPPEYNVCLGDLFSVAWLEDSDVQNSWYETLNQQYHHVDKRISHASHATQYGNL
KLGEEGLFVYMGSNPANDNYTSLDGNALTPSSIVVNQRDADLLHLWEKFRKAPEGSARKEEAQTQIFKAMSHRVHI
DSSIKLIGKLLFGIEKCTEILNAVRPAGQPLVDDWACLRSLVGTFETHCGSLSEYGMRHTRTIANICNAGISEEQMAEA
ASQACASIP*

pET21a-TP897-NGL-His$_6$:

MQVQLVECGGGLVQAGDSLRLSCVASGRSLDGATMRWYRQAPGKEREFVAGIFWDEIGTEYADTAKGRFTISRDNA
KNTIYLQMTNLRSEDTAMYYCNGLVFGGEYWGKGTLVTVSSNGLHHHHHH*

pET21a-TP1107-NGL-His$_6$:

MQVQLVECGGGLVQPGGSLRLSCAASGFTFSDTWMNWVRQAPGKGLYWISAINPDGGNTAYADSVKGRFTISRDN
AKNMVYLQMDNLRPEDTAMYYCAKGWVRLPDPDLVRGQGTQVTVSSNGLHHHHHH*

pET28a-His$_6$-EGFP:

MGSSHHHHHHMVSKGEELFTGVVPILVELDGDVNGHKFSVSGEGEGDATYGKLTLKFICTTGKLPVPWPTLVTTLTY
GVQCFSRYPDHMKQHDFFKSAMPEGYVQERTIFFKDDGNYKTRAEVKFEGDTLVNRIELKGIDFKEDGNILGHKLE
YNYNSHNVYIMADKQKNGIKVNFKIRHNIEDGSVQLADHYQQNTPIGDGPVLLPDNHYLSTQSALSKDPNEKRDH
MVLLEFVTAAGITLGMDELYK*

pET28a-His$_6$-mCherry:

MGSSHHHHHHMVSKGEEDNMAIIKEFMRFKVHMEGSVNGHEFEIEGEGEGRPYEGTQTAKLKVTKGGPLPFAWDIL
SPQFMYGSKAYVKHPADIPDYLKLSFPEGFKWERVMNFEDGGVVTVTQDSSLQDGEFIYKVKLRGTNFPSDGPVMQ
KKTMGWEASSERMYPEDGALKGEIKQRLKLKDGGHYDAEVKTTYKAKKPVQLPGAYNVNIKLDITSHNEDYTIVE
QYERAEGRHSTGGMDELYK*

$P_{Tight}$ (Tight TRE promoter)-EGFP-*PGK*-rtTA:

MVSKGEELFTGVVPILVELDGDVNGHKFSVSGEGEGDATYGKLTLKFICTTGKLPVPWPTLVTTLTYGVQCFSRYPD
HMKQHDFFKSAMPEGYVQERTIFFKDDGNYKTRAEVKFEGDTLVNRIELKGIDFKEDGNILGHKLEYNYNSHNVYI
MADKQKNGIKVNFKIRHNIEDGSVQLADHYQQNTPIGDGPVLLPDNHYLSTQSALSKDPNEKRDHMVLLEFVTAAG
ITLGMDELYK*
MSRLDKSKVINGALELLNGVGIEGLTTRKLAQKLGVEQPTLYWHVKNKRALLDALPIEMLDRHHTHFCPLEGESWQ
DFLRNNAKSFRCALLSHRDGAKVHLGTRPTEKQYETLENQLAFLCQQGFSLENALYALSAVGHFTLGCVLEEQEHQ
VAKEERETPTTDSMPPLLRQAIELFDRQGAEPAFLFGLELIICGLEKQLKCESGGPADALDDFDLDMLPADALDDFDL
DMLPADALDDFDLDMLPG*

*3×CRE*-mCherry-PEST:

MVSKGEEDNMAIIKEFMRFKVHMEGSVNGHEFEIEGEGEGRPYEGTQTAKLKVTKGGPLPFAWDILSPQFMYGSKA
YVKHPADIPDYLKLSFPEGFKWERVMNFEDGGVVTVTQDSSLQDGEFIYKVKLRGTNFPSDGPVMQKKTMGWEAS
SERMYPEDGALKGEIKQRLKLKDGGHYDAEVKTTYKAKKPVQLPGAYNVNIKLDITSHNEDYTIVEQYERAEGRHS
TGGMDELYKLSHGFPPEVEEQAAGTLPMSCAQESGMDRHPAACASARINV*

### Supplementary File 5. Primers for qPCR-based BLISA.

| **DNA barcode** | **primer** | **Sequence (5' to 3')** |
|---|---|---|
| **qbc. 1 (for qPCR)** | Forward | TCTTGTGGAAAGGACGAAACACG |
| | Reverse | CTAAAGCGCATGCTCCAGAC |
| **qbc. 2 (for qPCR)** | Forward | TACACGACGCTCTTCCGATC |
| | Reverse | ACTCGGTGCCACTTTTTCAA |
| **qbc. 3 (for qPCR)** | Forward | ACACGTCTGAACTCCAGTCAC |
| | Reverse | CAAGCAGAAGACGGCATACG |
| **qbc. 4 (for qPCR)** | Forward | GACAGTTCGAGTTTGAAGCGC |
| | Reverse | AGTATGCACTCCCACGTCTAG |
| **qbc. 5 (for qPCR)** | Forward | GAAAGATCTGGCTGCCATGC |
| | Reverse | TGATTCCAACCAGGTTTGCGA |
| **qbc. 6 (for qPCR)** | Forward | AGATGACGTCGATTGTTGGTCG |
| | Reverse | TACGGTGACACAACCTCCATG |
| **qbc. 7 (for qPCR)** | Forward | TCAGGTGCATAGGAGTCAGC |
| | Reverse | GGAGCCATGAACTGACAGCAT |
