## Supplementary material for "Modular DNA Barcoding of Nanobodies Enables Multiplexed *in situ* Protein Imaging and High-throughput Biomolecule Detection": Key Resources Table

| Key Resources Table | | | | |
|---|---|---|---|---|
| **Reagent type (species) or resource** | **Designation** | **Source or reference** | **Identifiers** | **Additional information** |
| strain, strain background (*Escherichia coli*) | SHuffle | Weidibio | EC2031 | |
| strain, strain background (*Escherichia coli*) | BL21 (DE3) | Transgene | CD601 | |
| strain, strain background (Mus musculus) | C57BL/6N | Beijing Vital River | 213 | |
| cell line (*Homo-sapiens*) | HEK293T | ATCC | CRL-3216 | |
| cell line (*Homo-sapiens*) | Hela | ATCC | CCL-2 | |
| cell line (*Homo-sapiens*) | U87 | ATCC | HTB-14 | |
| cell line (*Homo-sapiens*) | U937 | ATCC | CRL-1593.2 | |
| chemical compound, drug | DBCO-PEG$_3$-SS-NHS | Conju-Probe | CP-2089 | |
| chemical compound, drug | Anhydrotetracycline hydrochloride (ATc) | J&K | 541642 | |
| chemical compound, drug | Forskolin (FSK) | Sigma-Aldrich | F3917 | |

| | | | | |
|---|---|---|---|---|
| chemical compound, drug | 3-Isobutyl-1-methylxanthine (IBMX) | Sigma-Aldrich | I5879 | |
| chemical compound, drug | Anisomycin | MedChem Express | CS-4981 | |
| chemical compound, drug | Compounds in HTS-based BLISA | other | | See Supplementary File 1 |
| transfected construct | $P_{Tight}$-EGFP-*PGK*-rtTA | This paper | | See Supplementary File 4 |
| transfected construct | *3×CRE*-mCherry-PEST | This paper | | See Supplementary File 4 |
| biological sample (*Homo-sapiens*) | Human healthy (male) and psoriasis (female) skins | other | | See Materials and Methods |
| biological sample (*Homo-sapiens*) | Human serum | Beijing, China | | See Materials and Methods |
| antibody | anti-PDGFRα (Rabbit monoclonal) | Abcam | ab203491 | 1:500 |
| antibody | anti-KRT14 (Rabbit monoclonal) | other | | From Ting Chen's Lab; 1:1000 |
| antibody | anti-DCT (Rabbit monoclonal) | other | | From Ting Chen's Lab; 1:500 |
| antibody | anti-CD31 (Mouse monoclonal) | Abcam | ab9498 | 1:500 |
| antibody | anti-αSMA (Rabbit monoclonal) | Abcam | ab124964 | 1:500 |

| | | | | |
|---|---|---|---|---|
| antibody | anti-CD45 (Mouse monoclonal) | BD Biosciences | 555481 | 1:300 |
| antibody | anti-MAP2 (Rabbit monoclonal) | Thermo Fisher Scientific | PA5-85755 | 1:500 |
| antibody | anti-NeuN (Rabbit monoclonal) | Abcam | ab177487 | 1:500 |
| antibody | anti-TPH2 (Rabbit monoclonal) | Abcam | ab184505 | 1:200 |
| antibody | anti-TH (Rabbit monoclonal) | Merck Millipore | AB152 | 1:500 |
| antibody | anti-nNOS (Rabbit monoclonal) | Sigma-Aldrich | N7280 | 1:500 |
| antibody | anti-GFAP (Rabbit monoclonal) | Abcam | ab7260 | 1:750 |
| antibody | anti-Iba1 (Rabbit monoclonal) | FUJIFILM | 019-19741 | 1:500 |
| antibody | anti-DDC (Rabbit monoclonal) | Abcam | ab3905 | 1:50 |
| antibody | anti-NF-H (Rabbit monoclonal) | Thermo Fisher Scientific | 711025 | 1:100 |
| antibody | anti-TMEM119 (Rabbit monoclonal) | Abcam | ab209064 | 1:500 |
| antibody | anti-GABA (Rabbit monoclonal) | Sigma-Aldrich | A2025 | 1:750 |

| | | | | |
|---|---|---|---|---|
| antibody | anti-Orexin A (Rabbit monoclonal) | Phoenix Pharmaceuticals | H-003-30 | 1:500 |
| antibody | anti-5-HT (Rabbit monoclonal) | Immunostar | 20080 | 1:500 |
| antibody | anti-α-tubulin (Mouse monoclonal) | Sigma-Aldrich | T5168 | 1:40000 |
| antibody | anti-GFP (Rabbit polyclonal) | Thermo Fisher Scientific | A-11122 | 1:1000 |
| antibody | anti-mCherry (Rabbit polyclonal) | Thermo Fisher Scientific | PA5-34974 | 1:2000 for purified mCherry, 1:10000 for the expressed mCherry |
| antibody | anti-Human IgG (Rabbit monoclonal) | Abcam | ab181236 | 0.33 nM |
| antibody | anti-HBsAg (capture) (Goat monoclonal) | Beijing Wantai Biological | YTX2101 | 5 µg/mL |
| antibody | anti-HBsAg (detection) (Mouse monoclonal) | Beijing Wantai Biological | HBs-2C1 | 0.33 nM |
| antibody | anti-HBeAg (capture) (Mouse monoclonal) | Beijing Wantai Biological | 13B12-1 | 5 µg/mL |
| antibody | anti-HBeAg (detection) (Mouse monoclonal) | Beijing Wantai Biological | 9A4-1 | 0.033nM |
| antibody | anti-phospho-p38α (T180/Y182) (capture) (Mouse monoclonal) | R&D Systems | DYC869B | The vendor recommended conc. |

| | | | | |
|---|---|---|---|---|
| antibody | anti-phospho-p38α (T180/Y182) (detection) (Rabbit monoclonal) | R&D Systems | DYC869B | 1/10 of the vendor recommended conc. |
| antibody | anti-phospho-ERK1 (T202/Y204)/ERK2 (T185/Y187) (capture) (Mouse monoclonal) | R&D Systems | DYC1018B | The vendor recommended conc. |
| antibody | anti-phospho-ERK1 (T202/Y204)/ERK2 (T185/Y187) (detection) (Rabbit monoclonal) | R&D Systems | DYC1018B | 1/10 of the vendor recommended conc. |
| antibody | anti-phospho-JNK Pan Specific (capture) (Mouse monoclonal) | R&D Systems | DYC1387B | The vendor recommended conc. |
| antibody | anti-phospho-JNK Pan Specific (detection) (Rabbit monoclonal) | R&D Systems | DYC1387B | 1/10 of the vendor recommended conc. |
| antibody | anti-phospho-AMPKα1 (T183) (capture) (Goat monoclonal) | R&D Systems | DYC3528 | The vendor recommended conc. |
| antibody | anti-phospho-AMPKα1 (T183) (detection) (Rabbit monoclonal) | R&D Systems | DYC3528 | 1/10 of the vendor recommended conc. |
| antibody | anti-phospho-CREB (S133) (capture) (Goat monoclonal) | R&D Systems | DYC2510 | The vendor recommended conc. |
| antibody | anti-phospho-CREB (S133) (detection) (Rabbit monoclonal) | R&D Systems | DYC2510 | 1/10 of the vendor recommended conc. |
| antibody | anti-phospho-Src (Y419) (capture) (Goat monoclonal) | R&D Systems | DYC2685 | The vendor recommended conc. |

| | | | | |
|---|---|---|---|---|
| antibody | anti-phospho-Src (Y419) (detection) (Rabbit monoclonal) | R&D Systems | DYC2685 | 1/10 of the vendor recommended conc. |
| antibody | anti-phospho-Akt (S473) (capture) (Rabbit monoclonal) | Abcam | ab285034 | 6.0 μg/mL |
| antibody | anti-phospho-Akt (S473) (detection) (Rabbit monoclonal) | Abcam | ab285140 | 0.01 μg/mL |
| recombinant DNA reagent | pET28a-$His_6$-Ubiquitin-OaAEP1 (C247A) | PMID: 26680698 | | See Supplementary File 4 |
| recombinant DNA reagent | pET21a-TP897-NGL-$His_6$ | This paper | | See Supplementary File 4 |
| recombinant DNA reagent | pET21a-TP1107-NGL-$His_6$ | This paper | | See Supplementary File 4 |
| recombinant DNA reagent | pET28a-$His_6$-EGFP | This paper | | See Supplementary File 4 |
| recombinant DNA reagent | pET28a-$His_6$-mCherry | This paper | | See Supplementary File 4 |
| sequence-based reagent | HCR Initiators | PMID: 24712299 ; 38966983 | HCR probes | See Supplementary File 3 |
| sequence-based reagent | HCR Amplifiers | PMID: 24712299 ; 38966983 | HCR probes | See Supplementary File 3 |
| sequence-based reagent | DNA barcode oligos for qPCR | This paper | DNA barcodes | See Supplementary File 3 |
| sequence-based reagent | DNA barcode oligos for sequencing | This paper | DNA barcodes | See Supplementary File 3 |

| | | | | |
|---|---|---|---|---|
| sequence-based reagent | Primers for qPCR-based BLISA | This paper | qPCR primers | See Supplementary File 5 |
| peptide, recombinant protein | GVG-K($N_3$)-RG | Scilight-Peptide | | |
| peptide, recombinant protein | *Oa*AEP1 (C247A) | other | | See Materials and Methods |
| peptide, recombinant protein | TP897-NGL-$His_6$ | other | | See Materials and Methods |
| peptide, recombinant protein | TP1107-NGL-$His_6$ | other | | See Materials and Methods |
| peptide, recombinant protein | GFP | other | | See Materials and Methods |
| peptide, recombinant protein | mCherry | other | | See Materials and Methods |
| commercial assay or kit | BCA protein assay | Thermo Fisher Scientific | 23225 | |
| software, algorithm | Matlab | Mathworks | vR2018a | |
| software, algorithm | GraphPad Prism | GraphPad | v9.02 | |
| software, algorithm | Zen | Zeiss | v2.3 | |
| software, algorithm | LAS X | Leica | v5.1 | |

| | | | |
|---|---|---|---|
| software, algorithm | ImageJ | ImageJ | v 2.1.0 |
| software, algorithm | Python | Python | v3.8.10 |
| software, algorithm | UMI-tools | PMID: 28100584 | v1.1.2 |
| software, algorithm | Snakemake | PMID: 34035898 | v6.8.0 |
| software, algorithm | Biopython | PMID: 19304878 | v1.79 |
| software, algorithm | R | R | v4.0.3 |
| software, algorithm | RStudio | RStudio | v1.4.1103 |
| software, algorithm | Tidyverse | DOI: 10.21105/joss.01686 | |
